# Nucleosome positioning and sensitivity suggest novel functional organization of the purple sea urchin *(Strongylocentrotus purpuratus)* genome

**DOI:** 10.1101/2025.05.12.653534

**Authors:** Maya J. Munstermann, Jane M. Beniot, Sam E. Karelitz, Lorea N. Arambarri, Jonathan H. Dennis, Daniel K. Okamoto

**Affiliations:** Department of Integrative Biology, University of California Berkeley, Berkeley, CA 94720; Department of Biological Science, The Florida State University, Tallahassee, FL 32303

**Keywords:** nucleosome positioning, nucleosome sensitivity, purple sea urchin, chromatin architecture, gene regulation

## Abstract

The eukaryotic nucleosome is the fundamental subunit of chromatin and plays functional roles in DNA templated processes, including replication and transcription. In eukaryotic promoters, nucleosome organization is highly structured, with nucleosomes occupying canonical positions flanking the transcription start site (TSS), thereby regulating access of the transcriptional machinery to the underlying DNA. We determine whether this canonical distribution is present in the purple sea urchin, *Strongylocentrotus purpuratus*, a species of ecological importance and a model organism for developmental biology and climate science. We used titrations of micrococcal nuclease to produce high throughput maps of nucleosome distribution and sensitivity to digestion from female urchin gonad tissue. Unlike yeast, flies, zebrafish, maize, mice or humans, urchins have extended nucleosome repeat lengths and lack a nucleosome depleted region over TSSs. Urchin promoters are dominated by strongly positioned and highly occupied +1 and +2 nucleosomes which are most prominent in highly expressed genes. Additionally, urchin promoters exhibit distinct patterns of susceptibility to nuclease digestion, with heightened sensitivity upstream of the TSS and limited resistance to nuclease digestion. Discretely positioned sensitive nucleosomes were enriched in promoters of highly expressed genes, suggesting a relationship between nucleosome sensitivity and transcriptional regulation. Collectively, we present a comprehensive overview of the unique interplay between nucleosome positioning and chromatin sensitivity in sea urchins. Our study not only provides a better understanding of dynamics of gene expression in a key developmental organism, but also reveals potential heterogeneity in a key structural property of chromatin previously thought to be homogeneous in model eukaryotes.

**Graphical abstract:** 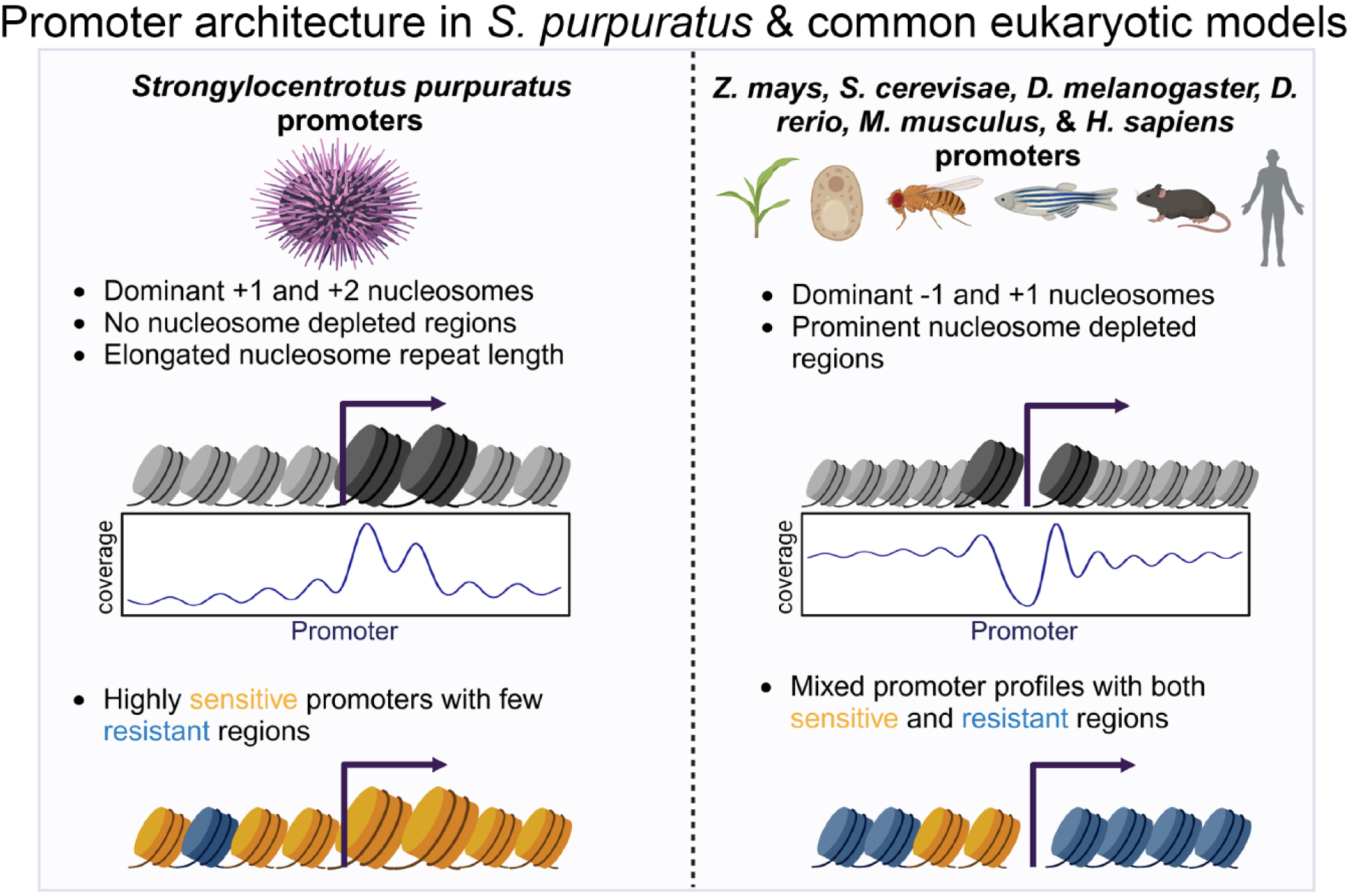

## Introduction

Eukaryotic chromosomes are composed of tightly packed chromatin which is the complex formed between proteins and DNA. Nucleosomes are the fundamental subunit of chromatin and are composed of ∼150bp of DNA that is wrapped 1.65 times around a core histone octamer. (Kornberg, 1974; Luger et al., 1997). Nucleosomes are organized along the DNA like beads on a string, with short stretches of linker DNA joining each nucleosome. These nucleosome arrays are further compacted into chromatin fibers which allows packaging of DNA into the nucleus. Nucleosomes have both structural and regulatory roles within eukaryotic genomes by packing DNA within the nucleus and regulating access to the underlying DNA (Kornberg and Lorch, 1999).

The organization of nucleosomes regulates access to the underlying DNA and thereby contributes to transcription, replication, repair, and any other DNA templated process (Kornberg & Lorch, 1999). Nucleosome distribution is especially important over transcription start sites (TSS) with densely packed nucleosomes preventing transcription factors and polymerases from accessing underlying DNA and transcribing genes (Bai & Morozov, 2010; Han & Grunstein, 1988; Jiang & Pugh, 2009; Yuan et al., 2005). Nucleosome repositioning or eviction from promoters can result in changes to structural features which allow for regulatory factor binding and recruitment of transcriptional machinery (Kornberg and Lorch 1999; Bell et al. 2011; Kaplan et al. 2009). Positioned nucleosomes can prevent the initiation of transcription by physically limiting assembly of the pre-initiation complex (Abril-Garrido et al., 2023; Lorch et al. 1987) and loss of nucleosomes can result in expression of formerly silent genes (Han & Grunstein, 1988). Elongating Polymerase II (pol II), the enzyme which transcribes DNA into mRNA, can fully displace nucleosomes (Lorch et al. 1987; Chen et al. 2019; Chang et al. 2013). Active promoters are marked by a pronounced nucleosome depleted region and a positioned +1 nucleosome, as shown in fruit flies, humans, and yeast (Gilchrist et al., 2010; Jimeno-González et al., 2015; Schones et al. 2008; Krebs et al. 2017; Jiang and Franklin Pugh 2009). Nucleosome positions are determined by sequence composition, barrier complexes, and chromatin remodeler activity (Chereji et al., 2018; Chereji & Clark, 2018; Gaffney et al., 2012; Gupta et al., 2008; Kornberg & Stryer, 1988). The position of nucleosomes around the TSSs regulates gene expression in a wide array of organisms (Bai & Morozov, 2010; Han & Grunstein, 1988; Jiang & Pugh, 2009; Schones et al., 2008; Weiner et al., 2015; Xi et al., 2011; Yuan et al., 2005, p. 200; Zhang et al., 2014).

Studies of chromatin structure have largely focused on the genomes of eukaryotic organisms frequently used in genetics research and have identified common features of nucleosome organization. Differences in tissue-specific (Clark et al., 2019), disease-specific (Druliner et al., 2016; Jacob et al., 2024; Piroeva et al., 2023), and species-specific nucleosome organization have been described (Tolstorukov et al., 2009) and show that nucleosome distribution is not static within or across well-studied organisms. Previous work has identified common features, such as nucleosome depleted regions, found over transcription start sites across a variety of taxa, including in archaea (Nalabothula et al., 2013), yeast (Lee et al., 2004; Mavrich et al., 2008b; Weiner et al., 2010; Yuan et al., 2005), fruit flies (Mavrich et al., 2008a; Mieczkowski et al., 2016), and humans (Kaplan et al., 2009; Schones et al., 2008). Additionally, nucleosomes tend to occupy discrete statistical positions that flank transcription start sites with gradual loss of phasing with increased distance from the promoter (Bai & Morozov, 2010; Kornberg, 1981; Kornberg & Stryer, 1988). However, this pattern of canonical or stereotypical nucleosome distribution over promoters is composed of discretely positioned dominant nucleosomes which together compose an average profile that generates the canonical pattern observed in bulk analysis (Cole et al., 2021; Cole & Dennis, 2020). While nucleosome distribution patterns are well understood in model eukaryotes, it is not clear how applicable these patterns are to other systems or non-model organisms.

Previous studies have shown that in addition to nucleosome positioning, nucleosome sensitivity also corresponds with gene expression and transcription factor binding in multiple organisms (Brahma & Henikoff, 2019; Cole et al., 2021; Mieczkowski et al., 2016; Parvathaneni et al., 2020; Pass et al., 2017; Rodgers-Melnick et al., 2016; Vera et al., 2014). To determine the locations of nucleosomes across the genome, Micrococcal nuclease (MNase) which can be used to cleave linker DNA allowing for high throughput mapping of nucleosome footprints (Schones et al., 2008; Shivaswamy et al., 2008; Sun et al., 1986). MNase digestions can be modified to include enzyme titrations to recover nucleosomes which are preferentially released under light or heavy digest conditions and identify regions that are sensitive or resistant to digestion, respectively (Clark et al., 2019; Cole et al., 2021; Cole & Dennis, 2020; Mieczkowski et al., 2016; Parvathaneni et al., 2020; Rodgers-Melnick et al., 2016; Vera et al., 2014; Weiner et al., 2010; Xi et al., 2011). Together, position and sensitivity profiles provide a comprehensive framework to disentangle chromatin structure and the associated gene regulatory mechanisms.

The purple sea urchin, *Strongylocentrotus purpuratus,* is an invertebrate deuterostome that is often used to model embryonic and larval developmental progression. While sea urchins were used for foundational studies of transcriptional regulation during development (Davidson et al., 2002; Halsell et al., 1987; Lieber et al., 1986; Mandl et al., 1997; Tu et al., 2012, 2014) and as a system for biochemical studies of nucleosome remodeling (Dong et al., 1990), little is known about genome wide chromatin structure. Sea urchins have hundreds of histone genes in clustered arrays which are temporally and spatially controlled during development (Cohn et al., 1976; Grunstein et al., 1981; Halsell et al., 1987; Kedes, 1979; Kedes et al., 1975; Lieber et al., 1986; Mandl et al., 1997; Maxson & Egzie, 1980). Additionally, embryonic urchin histone variants are known to alter nucleosome stability and DNase I digestion (Simpson, 1981). Purple sea urchins have the longest known eukaryotic nucleosome repeat length (∼250bp) which has been observed in sperm and some, but not all, developmental stages (Spadafora et al., 1976; Thomas et al., 1986; Widom et al., 1985). In sea urchin sperm, the elongated N-terminal tail of the sea urchin histone H2B binds specifically to linker DNA to stabilize chromatin structure. This unique feature may contribute to differences in linker length between urchins and other invertebrate and vertebrate species (Hill & Thomas, 1990). Additional studies have shown distinct regions of open chromatin, as defined by ATACseq, over urchin promoters during embryonic development (Arenas-Mena & Akin, 2023). Further, urchins are reported to have unique patterns of promoter-proximal Pol II pausing at the blastula stage (Chivu et al., 2023). These features suggest that urchin chromatin structure may differ from other eukaryotes. Current studies of urchin chromatin have focused on sperm and developing embryos and it remains unclear how nucleosomes are organized across the adult purple urchin genome.

To address this gap in knowledge, we comprehensively mapped nucleosome positioning and sensitivity to MNase in adult female purple sea urchin gonad tissue. We found that adult urchin ovaries have elongated linker DNA, lack a nucleosome depleted region, and have unusually strongly positioned and highly occupied +1 and +2 nucleosomes. The observed trends in urchin nucleosome distribution were not present in more commonly studied model organisms such as maize, yeast, fruit flies, zebrafish, mice, and humans. The novel organization of nucleosomes in urchins has important implications for understanding transcriptional modulation and regulation in a non-model organism as these patterns differ from any known eukaryote. We additionally found that urchin promoters are highly sensitive to MNase digestion and that distinct sensitivity patterns were associated with increased gene expression. Our results provide a framework for better understanding adult *S. purpuratus* chromatin architecture and methodology which can be adapted for non-model organisms. Understanding the nucleosome organization in purple sea urchins may reveal evolutionary variations in chromatin structure, genomic responses to environmental stressors, and gene regulation among eukaryotes.

## Results

We aim to understand nucleosome organization on a genome wide scale in adult female purple sea urchins. We show nucleosome organization by digesting crosslinked gonad tissue with a titration of micrococcal nuclease (MNase) to assay both nucleosome distribution and susceptibility to nuclease (described in Fig. 1). Sea urchin nucleosome distribution and nuclease susceptibility patterns are compared to those in maize (*Z. mays)*, yeast (*S. cerevisiae)*, fruit flies (*D. melanogaster)*, zebrafish (*D. rerio)*, mice (*M. musculus)*, and humans (*H. sapiens)*. These organisms are selected for comparative analysis as they are commonly used eukaryotic model organisms and represent highly divergent clades separated by millions of years of evolution (Fig. 1A). Previously published, publicly available MNase-seq data from each species was utilized for this comparative analysis and are processed in parallel with our urchin samples (Mieczkowski et al., 2016; Parvathaneni et al., 2020; Xi et al., 2011; Zhang et al., 2014). Briefly, nucleosomal fragments from heavy and light digests are used to determine the distribution of nucleosomes while the ratio of fragments from light versus heavy digests are used to determine nucleosome susceptibility to MNase digestion (Fig. 1B).

**Figure 1.**
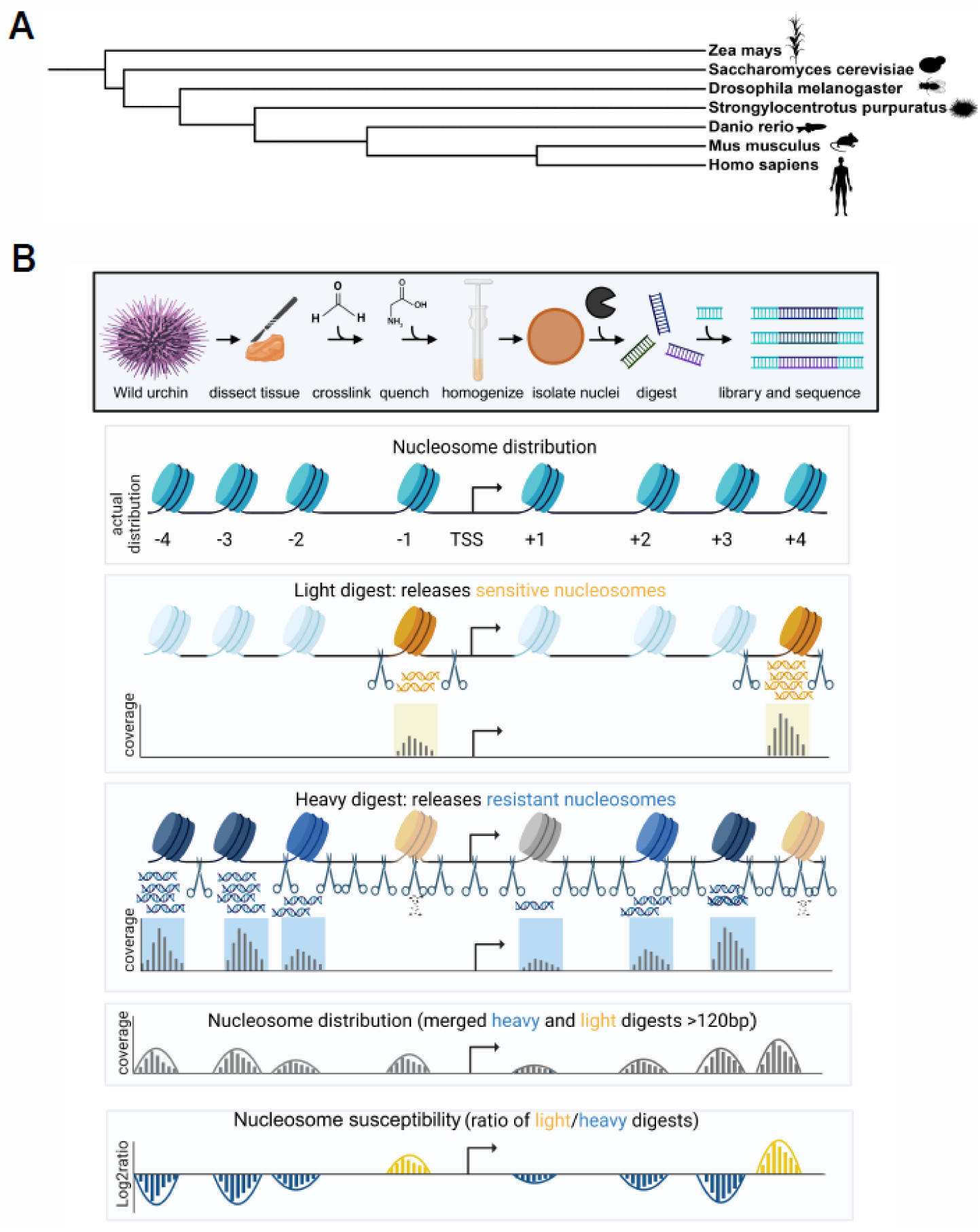
MNase based assays for probing chromatin architecture in sea urchins. A. Phylogenetic tree of model organisms used in comparative analyses. Branch lengths are not to scale. B. Schematic of nuclei isolation from urchin gonads and MNase digestion of crosslinked chromatin. Nuclei were digested with heavy and light MNase concentrations to determine nucleosome distribution and nuclease susceptibility. The combination of light and heavy digests was used to determine nucleosome distribution (grey). The log ratio of light over heavy digests was used to determine which nucleosomes were resistant (blue) or sensitive (gold) to MNase digestion.

### Nucleosome repeat length is elongated in *S. purpuratus* ovaries compared to other eukaryotic model organisms

We hypothesize that nucleosome characteristics would be similar in *S. purpuratus* compared to other eukaryotic model organisms considering that histone proteins are highly conserved in eukaryotes. However, MNase digestion of crosslinked urchin chromatin produced nucleosomal ladders with irregularly spaced nucleosomes compared to human samples which were processed in parallel (Fig. 2). Nucleosome ladders reveal the expected ∼150bp mononucleosome for both species. However, we also observe elongated linker DNA, as indicated by increased space between di-, tri-, and tetra-nucleosomes in urchin samples (Fig. 2A) compared to human samples (Fig. 2B). The urchin di-nucleosome is 400bp compared to 320bp for a human di-nucleosome. The urchin tri-nucleosome is 640bp compared to 490bp for the human counterpart, which reflects a nucleosome repeat length of ∼160bp for human cells compared to ∼250bp for urchin samples. Observed urchin nucleosome patterns are consistent in both light and heavy digest conditions (Supp. Fig. 1), suggesting that the extended linker length in sea urchins is an intrinsic feature of urchin chromatin rather than an enzyme titration effect. The extended linker length has previously been described for sea urchin sperm and developing embryos (Keichline & Wassarman, 1977; Spadafora et al., 1976; Thomas et al., 1986; Widom et al., 1985) with shorter linker lengths in the gastrula stage (Spadafora et al., 1976). The observed linker length in the ovary tissue is consistent with the dimer, trimer, and tetramer lengths previously described for different urchin tissues (Spadafora et al., 1976), indicating that this is a conserved feature of urchin chromatin in adult females ovaries as well as in urchin sperm and developing embryos.

**Figure 2.**
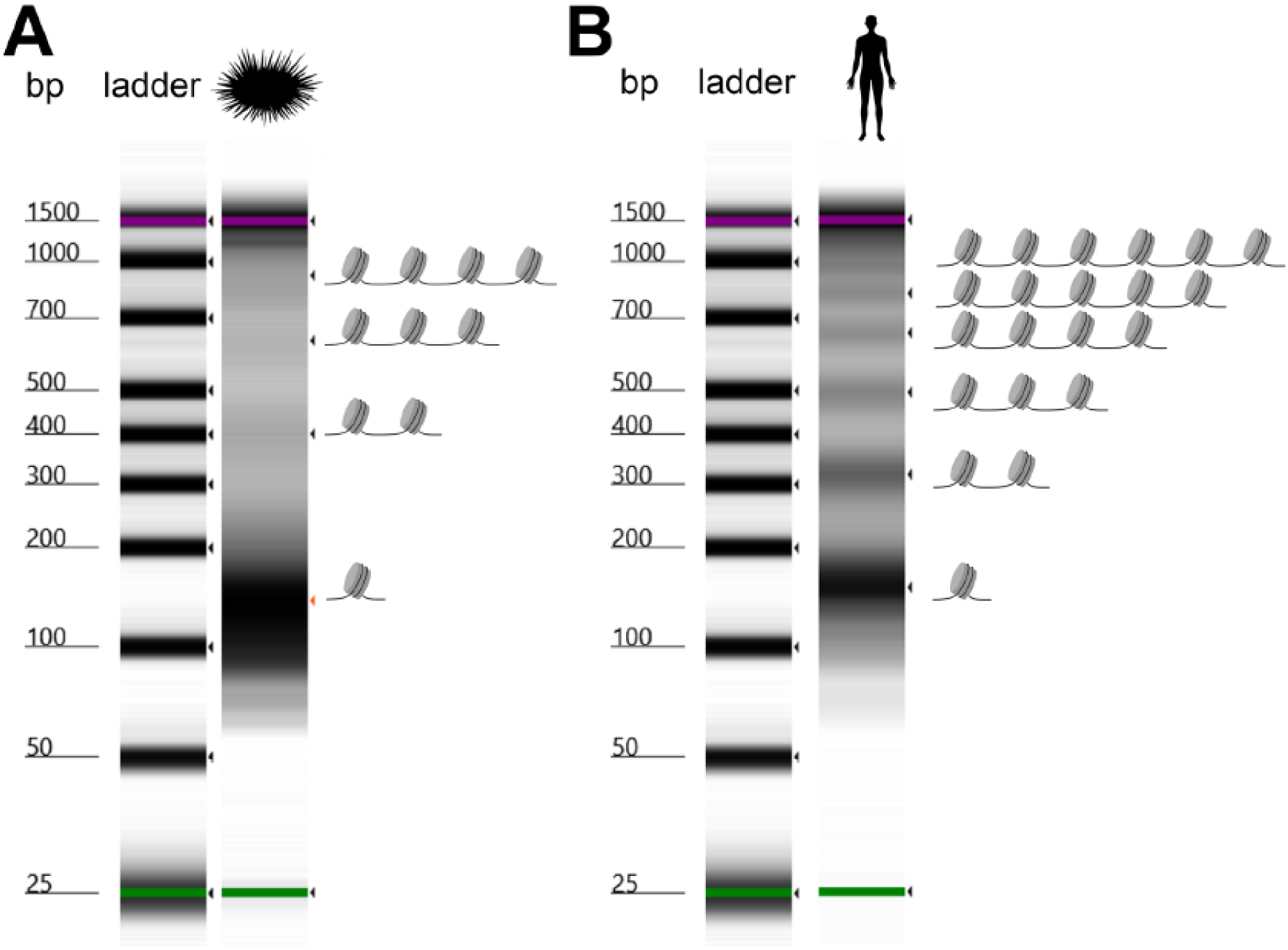
Sea urchins have elongated nucleosome repeat length compared to human samples. A. Representative tapestation gel of urchin gonad light digest with tick marks indicating mono-, di-, tri-, and larger nucleosome arrays. B. Representative tapestation gel of human THP-1 monocyte light digest with tick marks indicating mono-, di-, tri-, and larger nucleosome arrays.

**Supp. Fig. 1.**
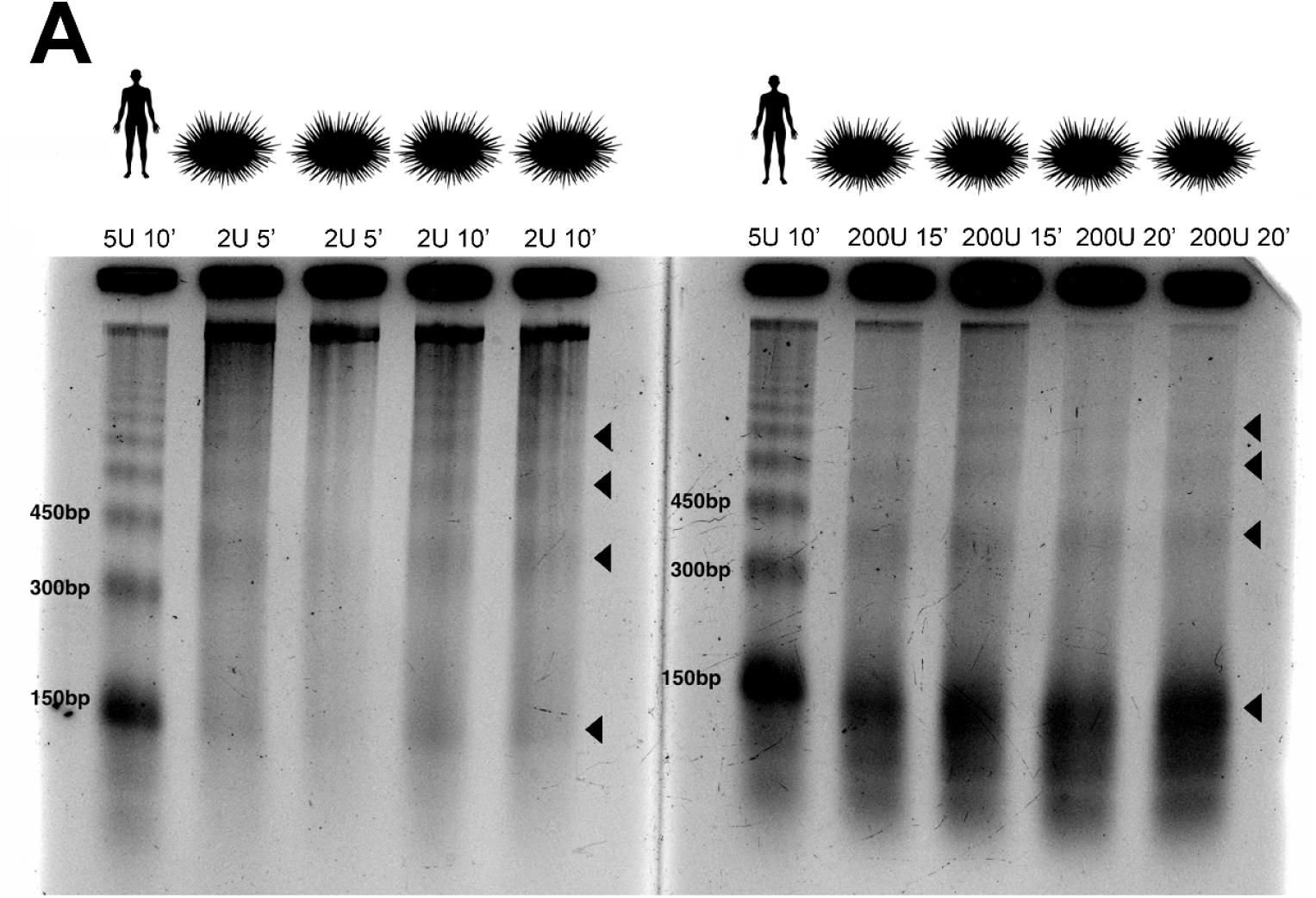
MNase digestion of crosslinked sea urchins gonad chromatin. A. Representative agarose gels of nucleosomal DNA isolated from heavy and light digests of urchin chromatin run beside a light digest of chromatin from human THP-1 cells. Sample names correspond to sample digest times and MNase concentrations. 2U 5’ samples were used as light digests and 200U 20’ were classified as heavy digests. Tick marks indicate urchin mono-, di-, tri-, and tetra-nucleosomes for both organisms.

### *S. purpuratus* promoters lack a defined nucleosome depleted region and are dominated by strongly positioned +1 and +2 nucleosomes

Next, we characterize nucleosome distribution patterns over sea urchin genes to compare with patterns observed in other eukaryotes. We compare nucleosome distribution over gene bodies in metagene space with 1kb flanking the transcription start site (TSS) and transcription end site (TES) (Fig 3A). Urchin genes are found to have well phased and highly occupied nucleosomes within promoters and moderate occupancy across the gene body with increasing signal at the TES. Poorly phased and weakly occupied nucleosomes are present beyond the TES (Fig. 3A). These patterns are consistent with previous observations in model organisms (Mavrich, et al., 2008b; Shivaswamy et al., 2008).

**Figure 3.**
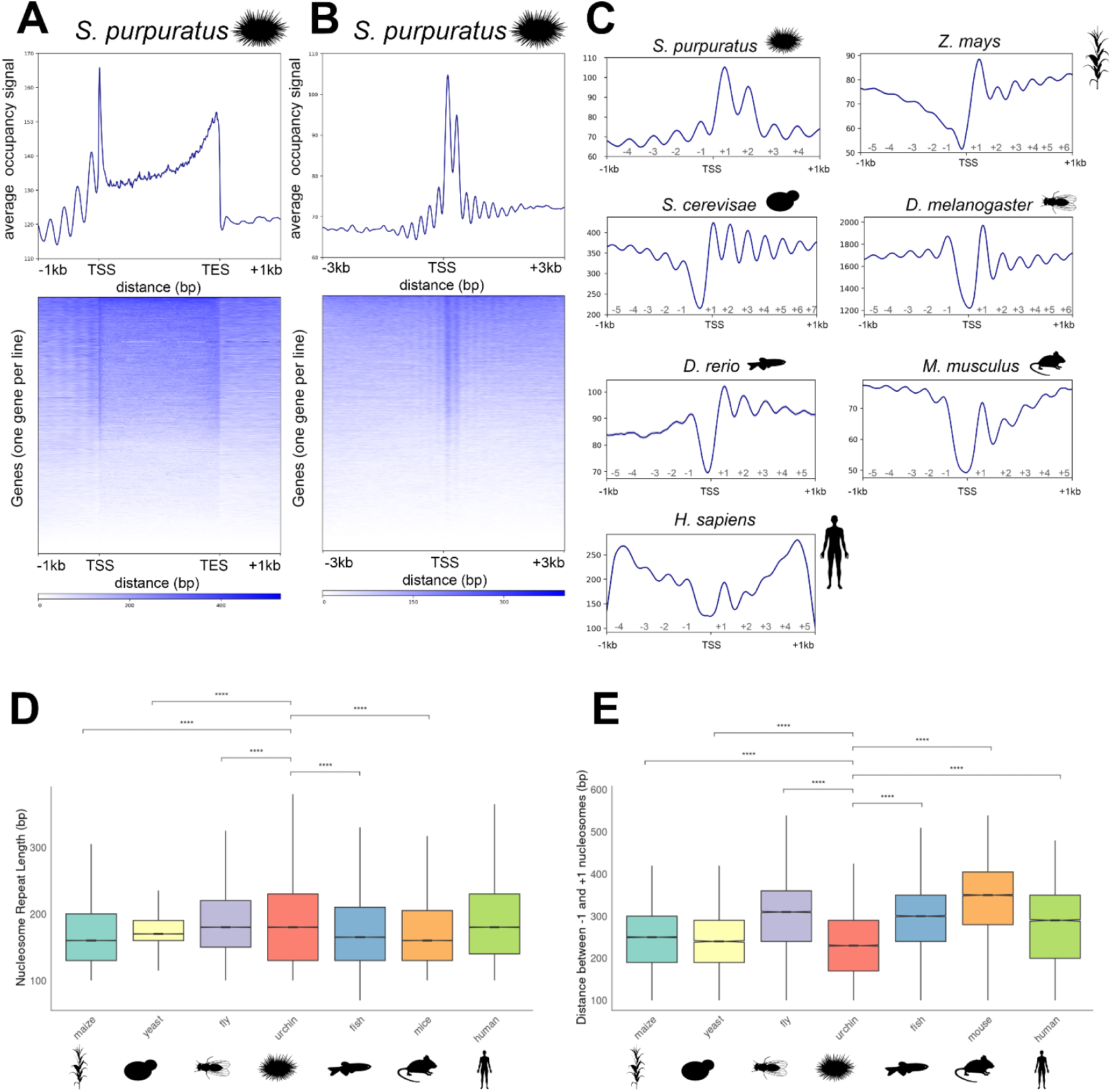
Sea urchin promoters lack a NDR and are dominated by strongly positioned +1 and +2 nucleosomes. A. Heatmap of nucleosome distribution over urchin genes in metagene space. The line plot above the heatmap shows the average distribution profile. Signal is shown over 1kb before the TSS over gene bodies in metagene space and 1kb beyond the TES. B. Heatmap of nucleosome distribution within 6kb of urchin TSSs. The average signal is shown in a line plot above the heatmap. C. Average nucleosome distribution profiles for urchin (*Strongylocentrotus purpuratus*), maize (*Zea mays*), yeast (*Saccharomyces cerevisiae*), fly (*Drosophila melanogaster*), zebrafish (*Danio rerio*), mouse (*Mus musculus*), and human (*Homo sapiens*) promoters centered on the TSS with 1kb flanking regions. Dominant nucleosomes are labeled in gray at the base of each plot. D. Nucleosome repeat length boxplots for all nucleosomes within 2kb of the TSSs of each model organism. Box plots represent the data in quartiles with the median shown as a notch, with the upper and lower whiskers representing Q1 and Q3. Asterisks indicate significant differences between urchin and other samples using the Kruskal-Wallis rank sum test. E. Boxplots showing the distance between the -1 and +1 nucleosomes of each model organism. Asterisks indicate significant differences (three asterisks: p <0.05, four asterisks: p <0.001) between urchin and other samples using the Kruskal-Wallis rank sum test.

We focus on nucleosome distribution on urchin promoters due to the direct role these nucleosomes play in gene regulation. We first examine nucleosome distribution over a 6kb window centered on the TSS and find that nucleosome occupancy and phasing are strongest over the immediate TSS with reduced phasing beyond 1kb upstream or downstream of the TSS (Fig. 3B). Because of this sharp loss of signal with increasing distance from the TSS, we define the promoter as 1kb upstream or downstream of the TSS for subsequent analyses. We validate these defined promoter regions by plotting publicly available *S. purpuratus* PROseq and Pol II ChIP data over these regions and we find significant enrichment in nascent RNA transcription and Pol II peaks over these regions, which confirm correct promoter annotation (Supp. Fig. 2).

**Supp. Fig. 2.**
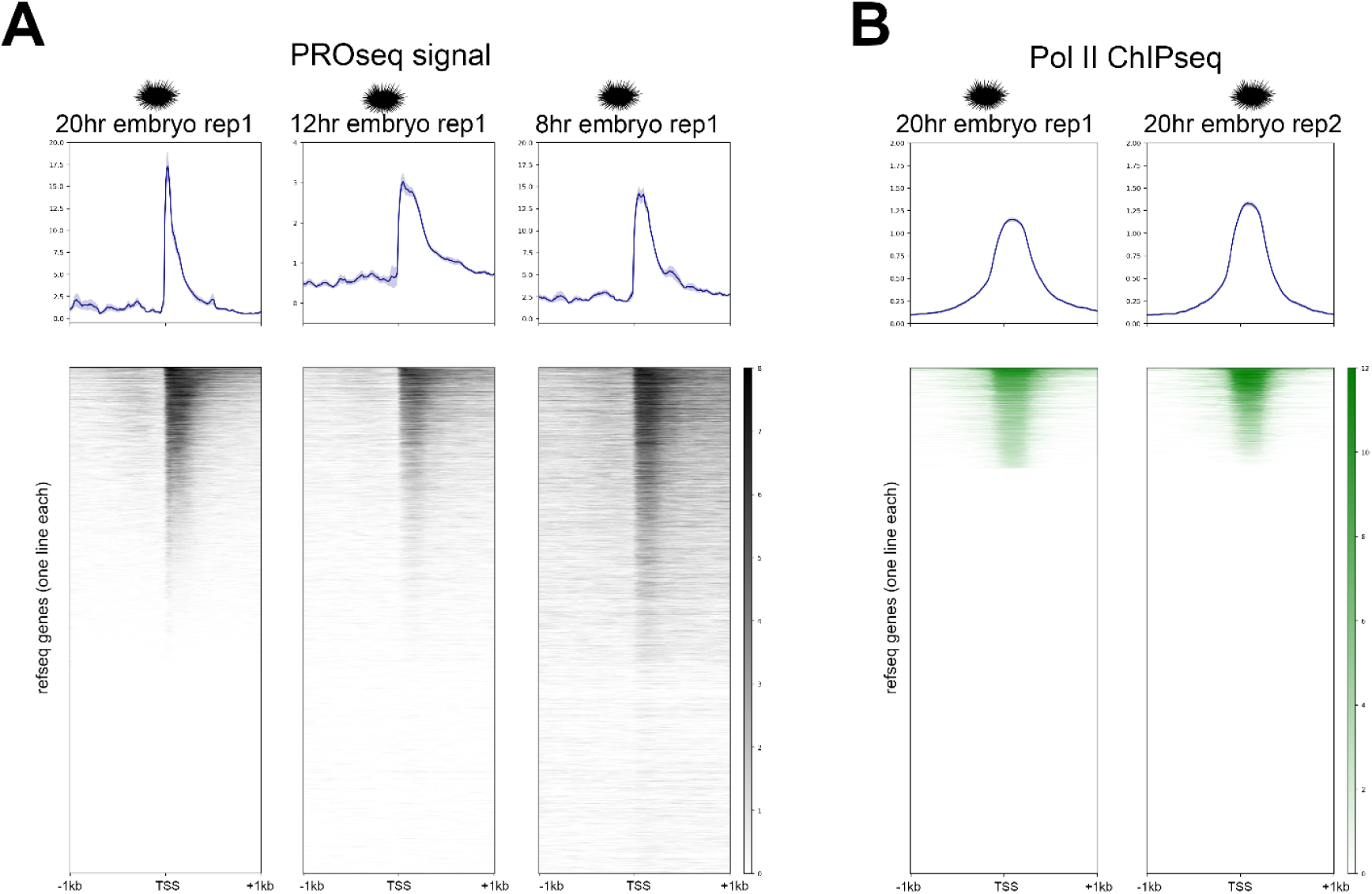
Annotated urchin promoters are enriched for nascent RNA transcription and polymerase II. A. Heatmaps show PROseq signal (black) over 2kb regions centered on annotated *S. purpuratus* TSSs. Signal is from developing embryos 20hr, 12hr, and 8hr post fertilization. B. Heatmaps show Pol II ChIPseq signal (green) over 2kb regions centered on annotated *S. purpuratus* TSSs. Signal is from developing embryos 20hr post fertilization.

With a closer analysis of nucleosome distribution over urchin promoters, we reveal regularly phased nucleosomes with especially high occupancy over the +1 and +2 nucleosome when promoters are viewed in aggregate (Fig. 3C). The nucleosome pattern over promoters is distinct from other eukaryotic model organisms. Firstly, urchin promoters lack a nucleosome depleted region (NDR) over the immediate TSS, which is well defined in the comparison of all model organisms (Fig. 3C). Secondly, urchin promoters have fewer nucleosomes within the defined 2kb promoter region due to longer linker length between nucleosomes (Fig. 3B). Despite lacking an NDR, urchins have 8 positioned nucleosomes within this window while other species have between 9-12 nucleosomes in the same spatially defined region. These features are consistent across multiple replicates of urchin tissue which were independently processed. These replicates show the same prominent +1 and +2 nucleosome patterns over promoter regions (Supp. Fig. 3). Within urchin promoters, nucleosomes are an average of 193.4bp (193.0-193.8 95% CI) apart which reflects a significantly longer distance compared to all other organisms apart from humans (Fig. 3D, Supp. Table 2). The 193bp linker length in urchin ovaries is shorter than the 250bp repeat length described for urchin sperm, potentially due to alternative histone variants expressed in sperm compared to ovaries (Lieber et al., 1986; Mandl et al., 1997; Simpson, 1981; Spadafora et al., 1976) or the location of histones within promoters compared to across the entire genome. Despite the increased nucleosome repeat length, urchins lack a well defined NDR, as observed in promoter maps (Fig. 3C) and by measuring the distance between the -1 and +1 nucleosome. This distance was significantly shorter in urchins compared to other species (Fig. 3E, Supp. Table 2). Notably, the difference between the distance between the -1 and +1 nucleosome and the mean nucleosome repeat length is considerably shorter in sea urchins at 39.6bp compared to all other organisms sampled, indicating little difference in spacing between these nucleosomes and any other nucleosome pair within the urchin promoter (Supp. Table 2). Together, the characteristic +1 and +2 nucleosomes and lack of an appreciable NDR mark urchin promoters as distinct from widely used model eukaryotes and suggest a novel mechanism for gene regulation in this species.

**Supp. Fig. 3.**
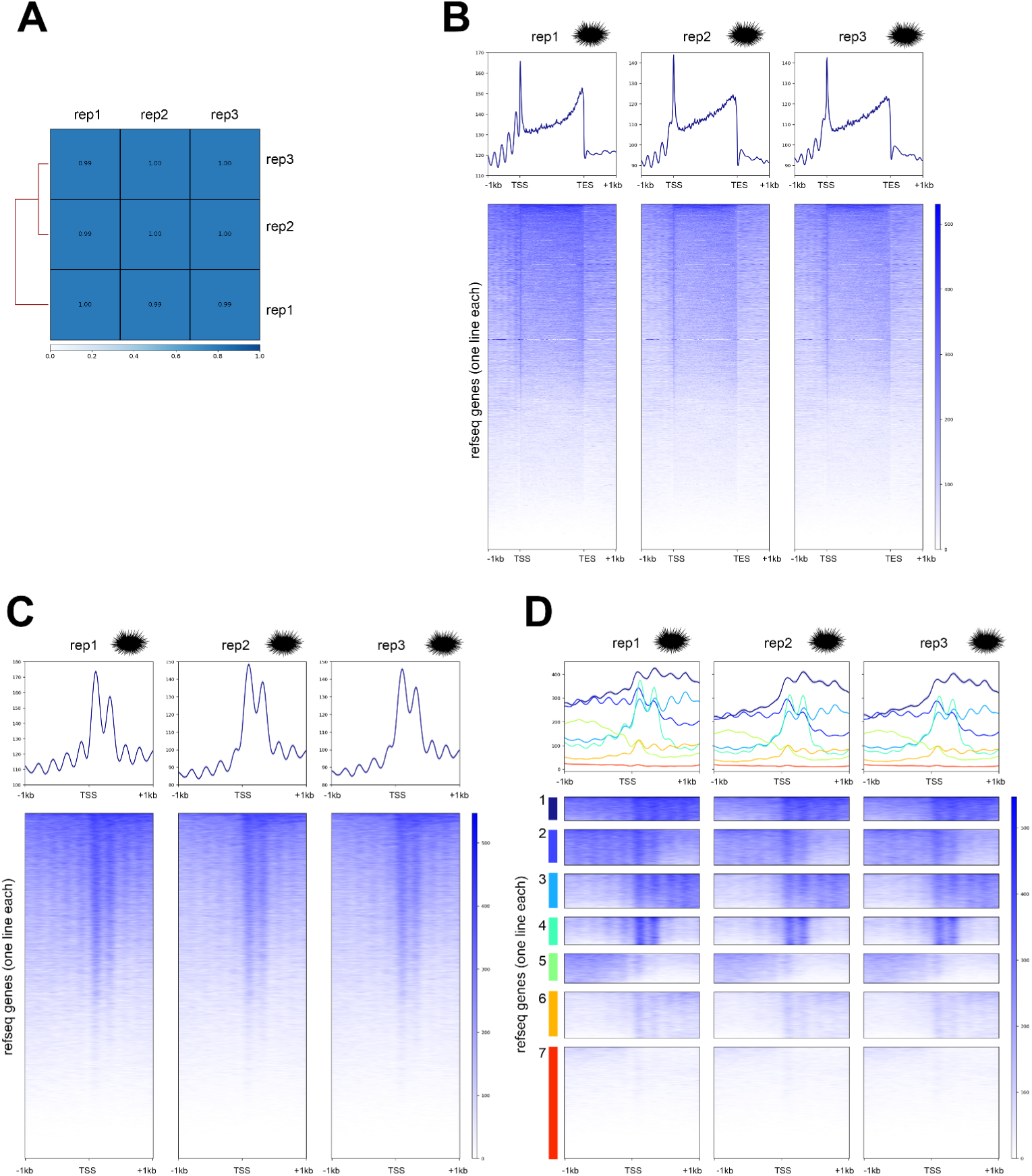
Nucleosome distribution is highly similar in replicate urchin tissue samples. A. Heatmaps of Pearson’s correlation of urchin distribution replicates. B. Heatmaps of nucleosome distribution (blue) over urchin genes in metagene space for each replicate sample. The line plot above the heatmap shows the average distribution profile for each replicate. C. Heatmaps of nucleosome distribution (blue) over urchin promoters in a 2kb window centered on the TSS. D. Kmeans clustered heatmaps showing nucleosome distribution (blue) over urchin promoters in a 2kb window centered on the TSS. Clusters are numbered and colored along the Y-axis of the heatmap.

**Supplemental Table 1.**
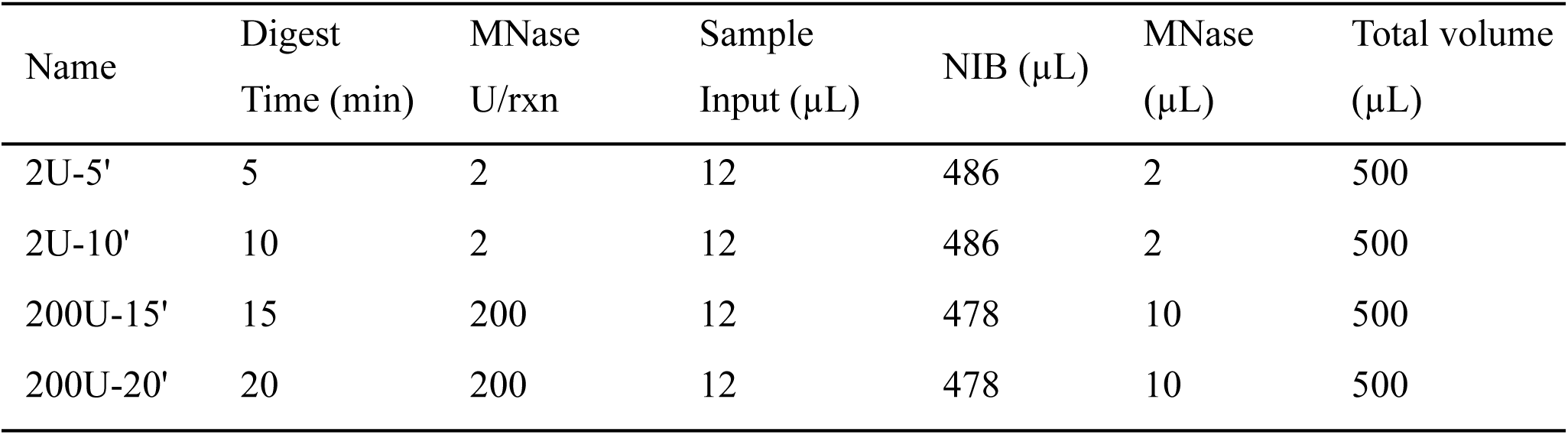
MNase digestion parameters for urchin gonad chromatin. MNase digestions parameters used for urchin chromatin. 2 unit (U) digests were combined for light digests and 200U digests were pooled for heavy digests. All digests were conducted at 37°C.

**Supplemental Table 2.** Sea urchins have elongated nucleosome repeat lengths and reduced nucleosome depleted regions compared to other model organisms. Mean + 95% confidence intervals (CI) for nucleosome repeat length within 2kb of TSSs in base pairs for model organisms. Mean + 95% confidence intervals for the distance between -1 and +1 nucleosomes in base pairs for model organisms. Differences between the distance between -1 and +1 nucleosomes and the mean nucleosome repeat length in bp.

| Organism | Mean nucleosome repeat length (bp) | Mean distance between -1 to +1 nucleosome (bp) | Difference between -1 to +1 distance vs mean repeat length (bp) |
| --- | --- | --- | --- |
| *Z. mays* | 174.6 (174.3-174.9 95% CI) | 249.9 (250.0 - 250.9 95% CI) | 75.3 |
| *S. cerevisiae* | 179.0 (178.6-179.5 95% CI) | 246.9 (245.3-248.5 95% CI) | 67.9 |
| *D. melanogaster* | 187.0 (186.7-187.3 95% CI) | 313.8 (311.9-315.6 95% CI) | 126.8 |
| *S. purpuratus* | 193.4 (193.0-193.8 95% CI) | 233.0 (231.9-234.0 95% CI) | 39.6 |
| *D. rerio* | 176.7 (176.3-177.0 95% CI) | 294.0 (292.5-295.4 95% CI) | 117.3 |
| *M. musculus* | 175.4 (175.2-175.6 95% CI) | 339.0 (337.6-340.5 95% CI) | 163.6 |
| *H. sapiens* | 195.2 (194.8-195.6 95%CI) | 277.1 (275.6-278.6 95%CI) | 81.9 |

### Strongly positioned +1 and +2 nucleosomes are unique to urchins and are found at the majority of active *S. purpuratus* genes

Due to the unique nucleosome features observed in sea urchin promoters, we want to determine if these patterns are generated by a small number of dominant genes or are common across the majority of promoters. To address this question, we generate nucleosome distribution maps for every RefSeq defined promoter in both urchins and other model organisms (Fig. 4A) and find that dominant features observed in the average plots were consistent across more than 60% of promoters in each organism (Fig. 4A). We then use kmeans clustering to group promoters by dominant nucleosome positions to resolve distinct promoter classes in each organism (Fig. 4B) and find that while each organism has variations in promoter classes, only the urchin samples show clusters defined by dominant +1 and +2 nucleosomes and lack an NDR (Fig. 4B). Examples of these patterns are seen over individual urchin promoters (Supp. Fig. 4). In total, 13,539 urchin promoters, representing 40% of all urchin promoters, have highly occupied +1 and +2 nucleosomes (Fig. 4B, clusters 1-4). These positioned +1 and +2 nucleosomes are also observed with lower signal values in the urchin promoter classes with lower overall occupancy (Fig. 4B, clusters 5-7). Other organisms show strongly positioned +1 or +2 nucleosomes but always in conjunction with an NDR (Fig. 4B, any non-urchin species). Clusters with poorly positioned or poorly occupied nucleosomes are observed in all organisms (Fig. 4B, cluster 7 all organisms) which may result from poorly positioned nucleosomes, highly repetitive sequences, or inaccessibility to enzymatic digestion. The unique promoter classes found in urchins further demonstrates that sea urchin ovaries have a promoter architecture which is distinct from other common model organisms.

**Supp. Fig. 4.**
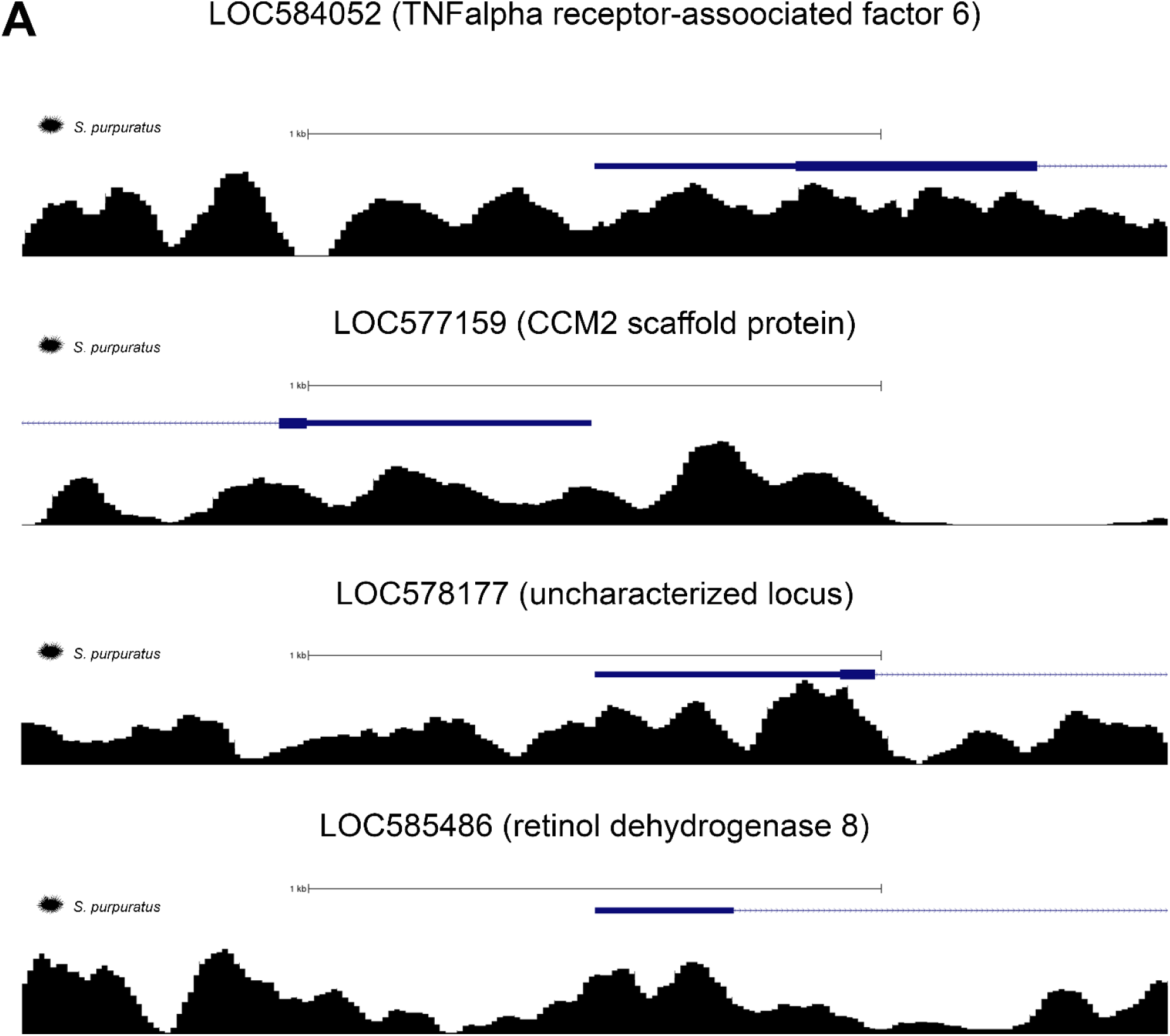
Nucleosome distribution over highly conserved genes is variable across species. Nucleosome distribution (black) over selected urchin gene promoters. Gene models (blue) for each species are shown above each distribution profile.

**Figure 4.**
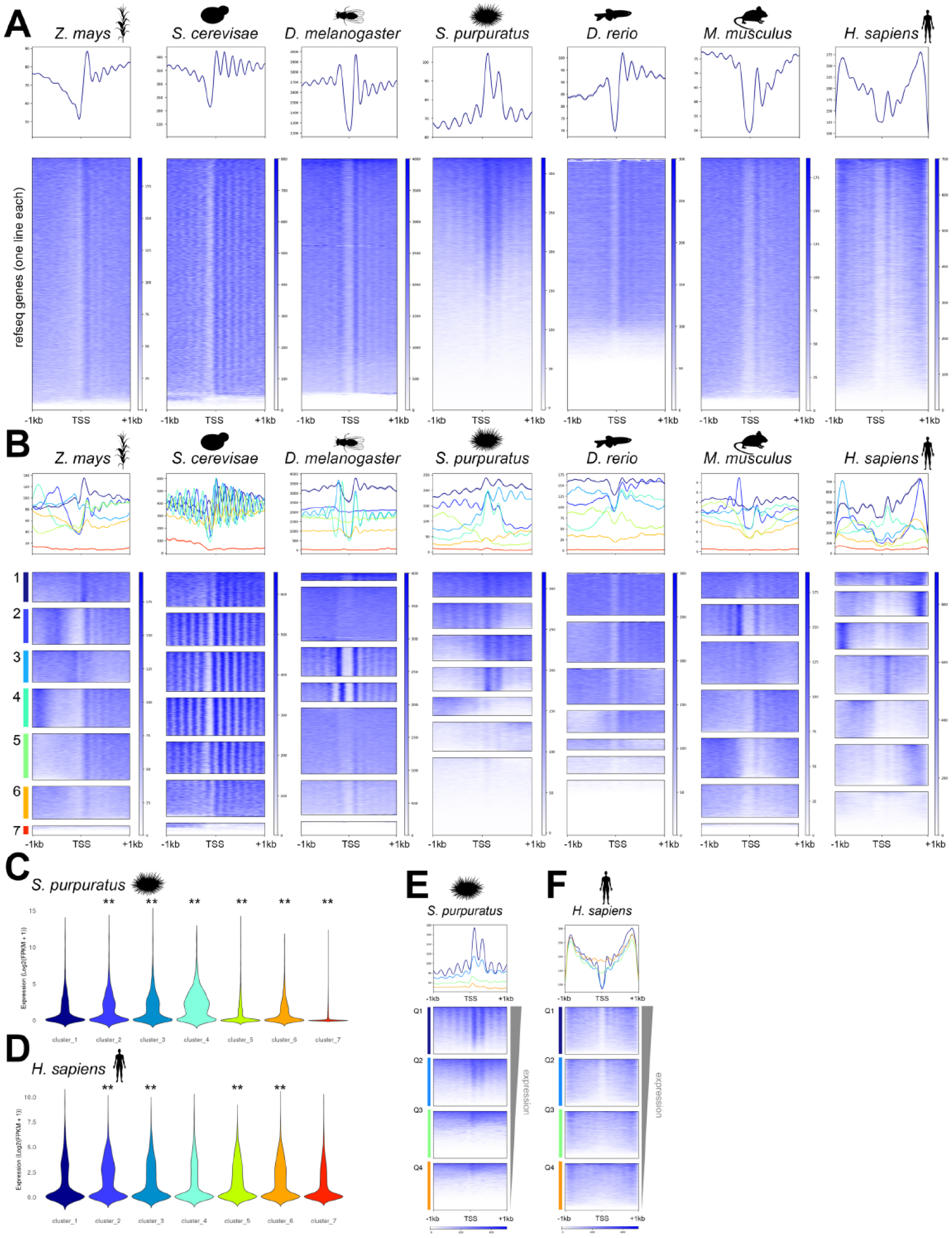
Dominant +1 and +2 nucleosomes are unique to urchins and are associated with elevated gene expression. A. Heatmaps showing nucleosome distribution (blue) over maize, yeast, fly, urchin, zebrafish, mouse, and human promoters in a 2kb window centered on the TSS. The line plots above the heatmap shows the average distribution profile for each organism. B. Independently generated kmeans clustered heatmaps showing nucleosome distribution (blue) over maize, yeast, fly, urchin, zebrafish, mouse, and human promoters in a 2kb window centered on the TSS. The line plot above the heatmap shows the average distribution profile for each cluster. Clusters are numbered and colored along the Y-axis of the heatmap. C. Urchin gene expression (log2(FPKM+1)) for genes within each kmeans cluster shown in B. Asterisks indicate significant differences (p-value < 0.0001) between cluster 1 and all other clusters using a Wilcoxon rank sum test. D. Human gene expression (log2(FPKM+1)) for each kmeans cluster shown in B. Asterisks indicate significant differences (p-value < 0.05) between cluster 1 and all other clusters using a Wilcoxon rank sum test. E. Urchin nucleosome distribution (blue) plotted over gene expression quartiles. F. Human nucleosome distribution (blue) plotted over gene expression quartiles.

Next, we determine whether the promoter class defined by highly occupied +1 and +2 nucleosomes correlates with gene expression. We hypothesize genes with this distribution pattern are poorly expressed as the presence of NDRs over the TSS are associated with higher gene expression model organisms (Bai & Morozov, 2010; Schones et al., 2008; Shivaswamy et al., 2008). To test this hypothesis, we compare publicly available urchin ovary gene expression data (Tu et al., 2012, 2014) with our nucleosome distribution clusters. We find that urchin promoters with highly occupied +1 and +2 nucleosomes lacking an NDR have significantly higher gene expression compared to other clusters (Kruskal-Wallis rank sum test p-value < 2.2e-16) (Fig. 4C, compare clusters 1-4 vs clusters 5-7). This observation is unexpected as highly occupied nucleosomes without a NDR are thought to inhibit transcriptional machinery from binding promoter regions. Analysis of human promoter clusters show that each promoter class had more similar expression levels with the exception of the poorly positioned cluster which has lower overall expression (Kruskal-Wallis rank sum test p-value < 2.2e-16) (Fig. 4D). We confirm our result by plotting nucleosome distribution over gene expression quartiles in both organisms. This analysis shows that the highest expressing urchin genes are dominated by strongly positioned +1 and +2 nucleosomes (Fig. 4E). This contrasts sharply with highly expressed human genes which are dominated by a pronounced NDR flanked by positioned nucleosomes (Fig. 4F). Both urchins and humans show an association between highly positioned nucleosomes and elevated gene expression, but the nature of these positioned nucleosomes varies considerably between species in terms of NDRs. The lack of an NDR in highly expressed urchin genes suggests that urchins may have a different mechanism for recruiting transcription factors and transcriptional machinery or nucleosome remodeling which is not observed in other model organisms.

### *S. purpuratus* promoters are highly sensitive to MNase digestion and exhibit unique sensitivity patterns

We next evaluated urchin susceptibility to MNase digestion to determine if nucleosome sensitivity also exhibited a novel pattern. To test this hypothesis, we calculated the log2ratio of light to heavy digests across the urchin genome and plotted resulting nucleosome sensitivity over urchin gene bodies flanked by 1kb regions (Fig. 5A). Urchin promoters and the region downstream of the TES are largely sensitive to MNase digestion while gene bodies have mixed sensitivity signals. To determine if common sensitivity patterns are found across urchin genes, we use a kmeans clustering algorithm to group urchin genes based on similar sensitivity features (Fig. 5B). The clustering analysis reveals four classes of genes with distinct sensitivity patterns:

1. three clusters with strongly positioned sensitive regions over promoters (Fig. 5B clusters 2-4),
2. one cluster with a strongly positioned resistant region over the promoter (Fig. 5B, clusters 6),
3. two clusters with dominant sensitive positions in the TES region (Fig. 5B, clusters 1 & 5), and
4. one cluster lacking dominant features (Fig. 5B, cluster 7). These patterns correspond with similar observations in other organisms wherein promoters tend to be especially sensitive to MNase digestion (Cole et al., 2021; Cole & Dennis, 2020; Mieczkowski et al., 2016; Rodgers-Melnick et al., 2016; Vera et al., 2014; Xi et al., 2011).

**Figure 5.**
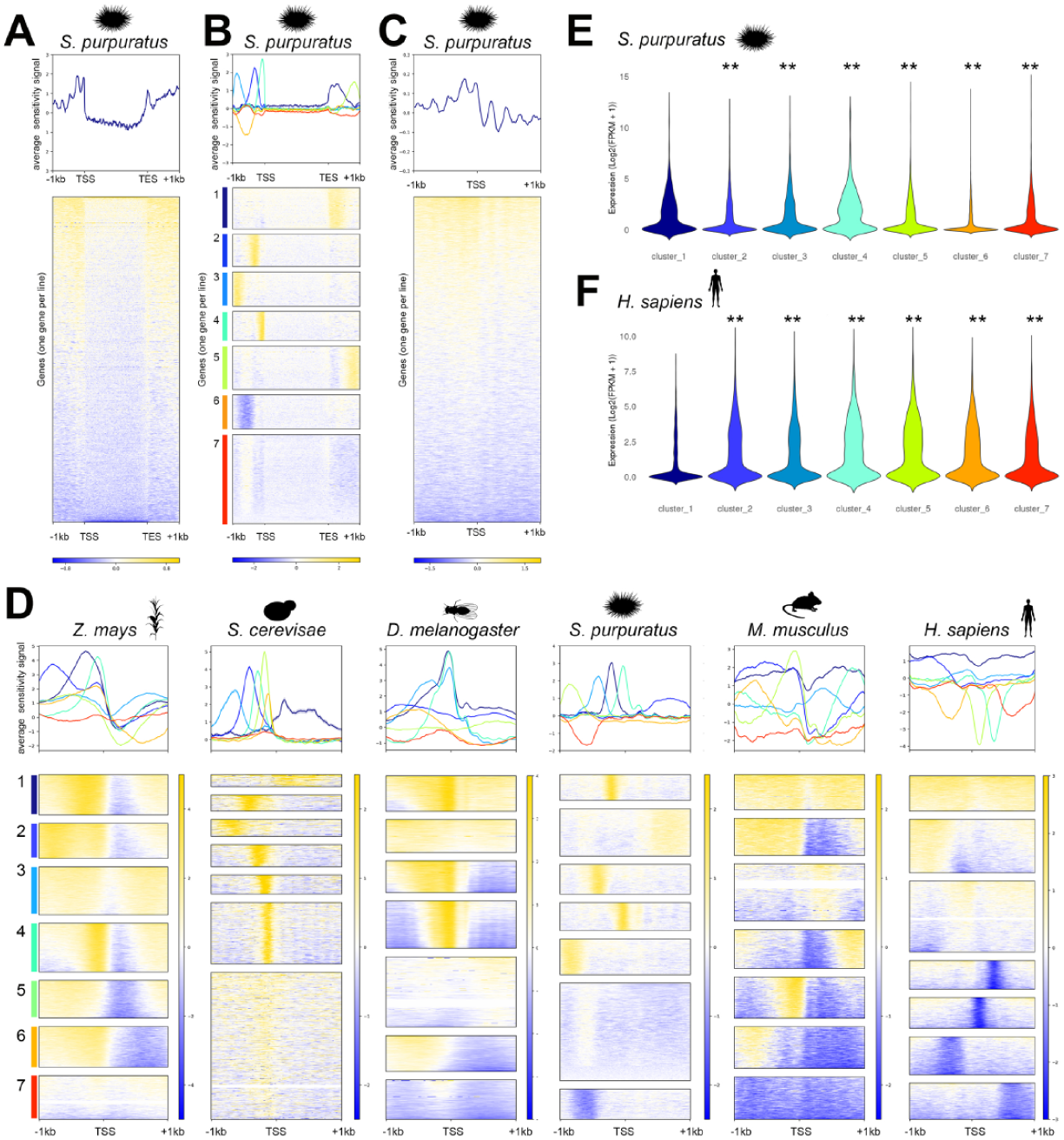
Highly sensitive urchin promoters are highly expressed. A. Heatmap of urchin nucleosome susceptibility over genes in metagene space with 1kb upstream of the TSS and 1kb downstream of the TES. Sensitive regions are shown in gold and resistant regions are shown in blue. A line plot above the heatmap shows the average susceptibility signal with positive values reflecting sensitive nucleosomes and negative values representing resistant nucleosomes. B. Kmeans clustering of the same susceptibility data over the same genes in metagene space as shown in A. Kmeans clusters are colored and labeled along the y axis. C. Heatmaps showing nucleosome sensitivity over a 2kb window centered on the TSS. D. Kmeans clustered heatmaps of nucleosome susceptibility over maize, yeast, fly, urchin, mouse, and human promoters. Clusters are colored and labeled along the y axis. E. Urchin gene expression (log2(FPKM+1)) for each susceptibility cluster shown in D. Asterisks indicate significant differences (p-value < 0.0001) between cluster 1 and all other clusters using a Wilcoxon rank sum test. F. Human gene expression (log2(FPKM+1)) for each susceptibility cluster shown in D. Asterisks indicate significant differences (p-value < 0.0001) between cluster 1 and all other clusters using a Wilcoxon rank sum test.

The greatest number of distinct susceptibility clusters are localized within promoters. Thus we focused on this region for all subsequent analyses. We first generated sensitivity maps over 2kb promoter regions flanking the TSS of all urchin genes (Fig. 5C). The resulting maps showed discretely positioned sensitive nucleosomes which are strongest over the -2 and -1 positions (Fig. 5C, note stripes in heatmap and periodicity in average plot). We use kmeans clustering of promoter sensitivity to clarify these patterns and generate groups of genes with similar promoter features in urchins and the other model organisms (Fig. 5D). The clustering analysis show five clusters of urchin genes with discretely positioned sensitivity patterns (Fig. 5D, urchin clusters 1-5), a mixed cluster that lacked defining features (Fig. 5D, urchin cluster 6), and a strongly resistant upstream cluster (Fig. 5D, cluster 7). While an additional upstream resistant cluster is noted in the second urchin replicate, the overall patterns were dominated by positioned sensitive nucleosomes upstream of the TSS (Supp. Fig. 5). The urchin sensitivity pattern is most similar to maize, yeast, and flies which were also dominated by sensitivity clusters which tend to be upstream of the TSS. However, only yeast produced the same type of discretely positioned downstream sensitivity patterns and broad upstream sensitivity cluster observed in urchins (Fig. 5D, compare *S. cerevisiae* and *S. purpuratus* average plots) despite urchins and yeast having vastly different gene structures, genome size, and intron distribution (Neuvéglise et al., 2011). The human and mouse samples show distinct promoter classes which are not observed in urchins or other organisms. These promoter classes included discreetly positioned resistant nucleosomes (e.g. human clusters 4-7) and diffusely sensitive patterns which spanned the entire promoter region (Fig. 5D, mouse and human cluster 1). Despite the differences seen in overall nuclease susceptibility patterns, highly conserved individual genes tend to be highly sensitive over the immediate TSS in all organisms (Supp. Fig. 6). This suggests that while overall sensitivity patterns may be distinct across species, there are conserved features which may underlie commonalities in chromatin structure across evolutionary time.

**Supp. Fig. 5.**
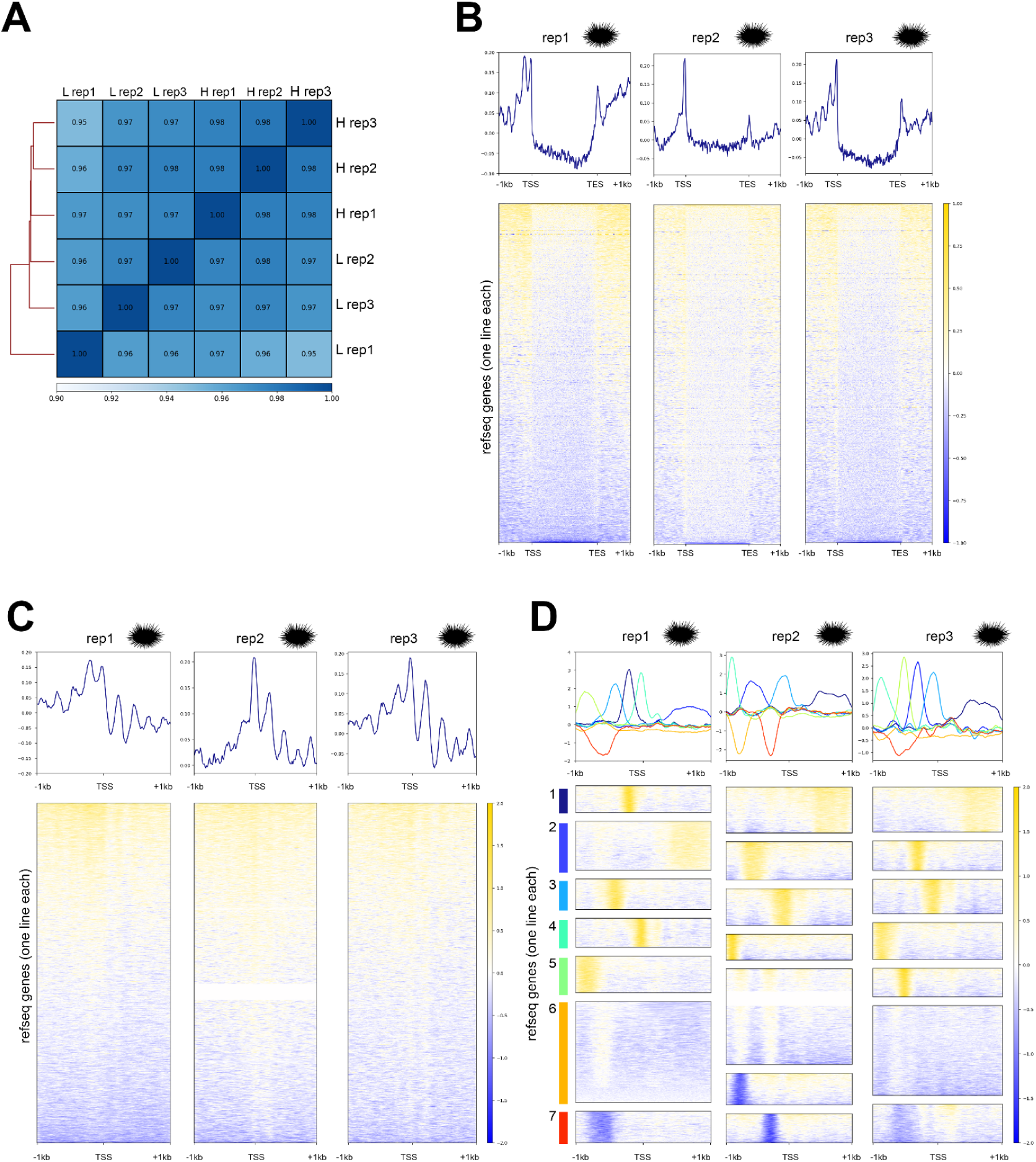
Susceptibility profiles are highly similar over replicate urchin samples. A. Heatmaps of Pearson’s correlation of urchin heavy (H) and light (L) replicates to show similarity proportions between replicate samples. B. Heatmaps showing nucleosome susceptibility over urchin genes in metagene space with 1kb upstream of the TSS, gene bodies, and 1kb downstream from the TES for each replicate sample. The line plot above the heatmap shows the average susceptibility profile for each replicate. C. Heatmaps showing nucleosome susceptibility over urchin promoters in a 2kb window centered on the TSS. The line plots above the heatmap shows the average susceptibility profile for each replicate. D. Kmeans clustered heatmaps showing nucleosome susceptibility over urchin promoters in a 2kb window centered on the TSS. The line plot above the heatmap shows the average profile for each cluster. Clusters are numbered and colored along the Y axis of the heatmap.

**Supp. Fig. 6.**
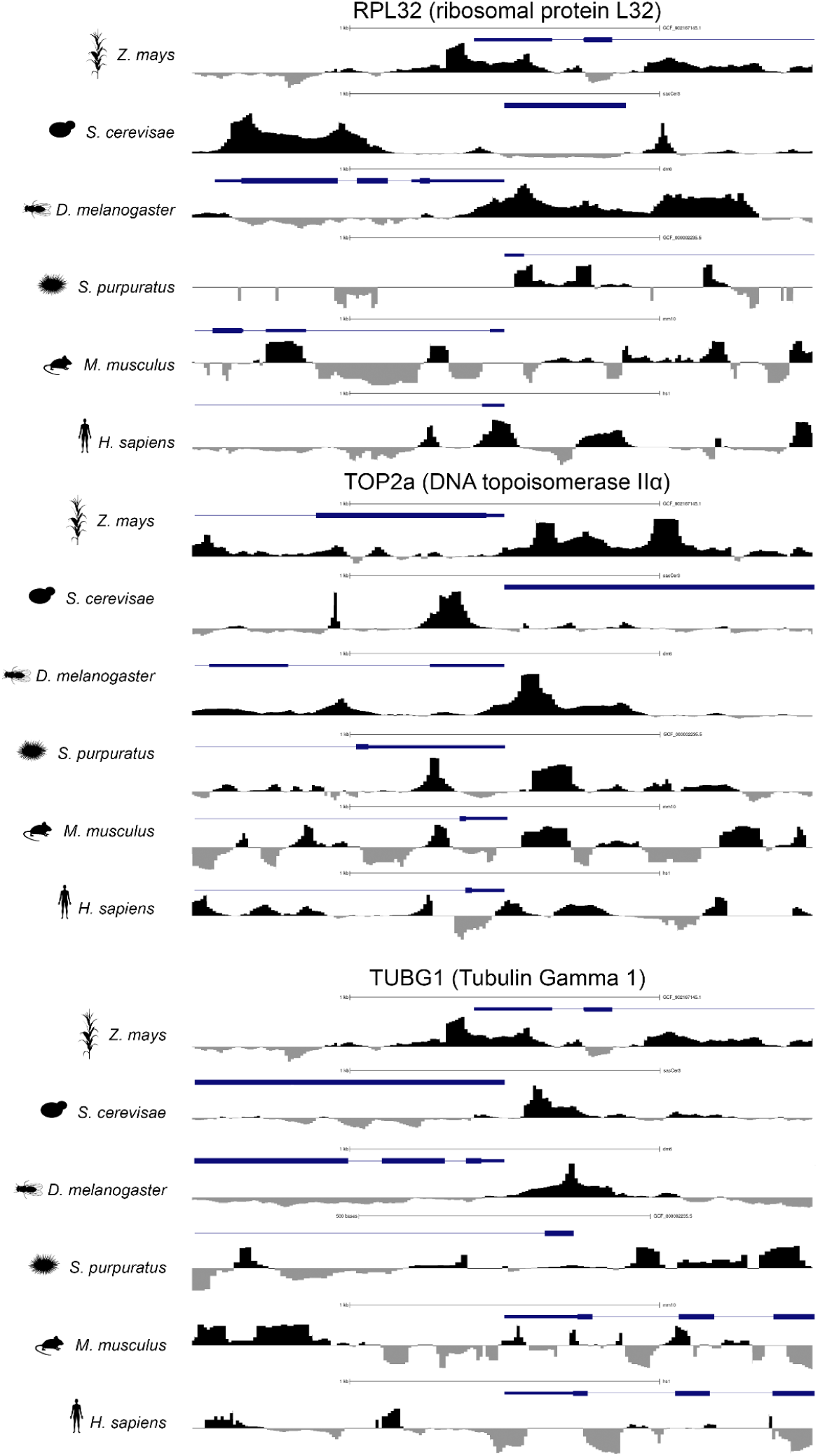
Nucleosome susceptibility over highly conserved genes. Nucleosome susceptibility for model organisms over RPL32, TOP2a, and TUBG1 gene promoters. Sensitive regions are shown in black while resistant regions are shown in gray. Gene models (blue) for each species are shown above each distribution profile.

In humans and maize, MNase sensitivity reveals promoters with discrete sensitive and resistant regions that are highly expressed (Cole et al., 2021; Cole & Dennis, 2020; Rodgers-Melnick et al., 2016; Vera et al., 2014). To determine if this trend is also observed in urchins, we plot urchin gene expression over each sensitivity cluster (defined in Fig. 5D). We find the two sensitivity clusters, defined by highly sensitive positioned nucleosomes just upstream or over the TSS (Fig. 5D, clusters 1&4), have significantly higher expression in urchins compared to other susceptibility clusters (Kruskal-Wallis rank sum test p-value < 2.2e-16) (Fig. 5E). The other sensitivity clusters have similar expression profiles as the positioned resistant cluster and the genes in the mixed cluster are poorly expressed (Fig. 5E). The relationship between urchin sensitivity clustering and gene expression in urchins contrasts with human cells, in that the greatest expression in human cells is associated with positioned resistant clusters and the diffusely sensitive cluster is largely silent (Kruskal-Wallis rank sum test p-value < 2.2e-16)(Fig. 5F). This suggests that both the location and intensity of sensitivity is associated with gene expression and that nuclease susceptibility profiles may have different relationships with gene regulation in sea urchins compared to mammals.

## Discussion

Despite the rich history of *Strongylocentrotus purpuratus’* use as a developmental model organism, and its ecological importance, the chromatin architecture and gene regulatory programs of adult organisms are poorly understood. In this study, we adapt genomic techniques to probe chromatin organization and sensitivity to enzymatic digestion in wild-caught adult female *S. purpuratus* gonad tissue to better understand relationships between nucleosome dynamics and gene expression in urchin ovaries. We successfully adapt MNase sequencing methodology for use with *S. purpuratus* gonad tissue by modifying crosslinking conditions and MNase digestions to achieve titrations suitable for nuclease susceptibility assays. The resulting nucleosome maps are the first high-throughput MNase-seq maps produced for adult *S. purpuratus* gonads and provide foundational insights into nucleosome distribution in this species. Importantly, our analysis centers on ovary tissue and it is possible that we included both adult gonad and oocyte tissue in our samples. However, multiple tissue samples produce highly consistent nucleosome distribution and sensitivity maps suggesting that if the sample is mixed, it is remarkably consistent across different replicates (Supp. Fig. 3 & Supp. Fig. 5).

We find that *S. purpuratus* gonad tissue has extended nucleosome repeat length compared to other eukaryotic organisms, which is aligns with previous work showing elongated linker DNA in urchin sperm and developing embryos (Keichline & Wassarman, 1977, 1979; Spadafora et al., 1976; Thomas et al., 1986; Widom et al., 1985). This result has implications for both gene regulation and higher order chromatin architecture as linker length facilitates nucleosome stacking and contributes to overall chromatin structure (Gibson et al., 2020; Szerlong & Hansen, 2011; Correll et al., 2012; Dias & D’Arcy, 2025) Extended linker length contributed to fewer nucleosomes within urchin promoters compared to other eukaryotic model organisms. Additionally, urchin promoters are characterized by strongly positioned, highly occupied +1 and +2 nucleosomes without a substantial nucleosome-depleted region (NDR), diverging from promoter architectures reported in other model organisms such as archaea, plants, yeast, flies, fish, mice, and humans (Bai & Morozov, 2010; Cole et al., 2021; Mieczkowski et al., 2016; Nalabothula et al., 2013; Parvathaneni et al., 2020; Xi et al., 2011; Yuan et al., 2005; Zhang et al., 2014). Interestingly, the lack of an NDR has also previously been described in giant virus genomes (Bryson et al., 2022) which use a novel chromatin packaging mechanism and lack linker DNA. Unique urchin histone variants are known to stabilize chromatin at different developmental stages (Grunstein et al., 1981; Halsell et al., 1987; Kedes, 1979; Leiber et al., 1986; Mandl et al., 1997; Marzluff et al., 2006; Maxson & Egzie, 1980; Green & Poccia, 1985; Hill et al., 1990; Hill & Thomas, 1990; Keichline & Wassarman, 1977; Poccia & Green, 1992; Simpson, 1981; Simpson & Bergman, 1980). These variants may contribute in additional stabilization of a nucleosome over the immediate TSS which is more fragile and prone to overdigestion in other organisms. If this is the case, the +1 and +2 nucleosomes are especially stable as robust signals are observed in both light and heavy digest conditions. Stabilization may be the result of a marine specific mechanism to resist salt induced nucleosome disassembly (Carruthers et al., 1998). Intriguingly, promoters with the highest +1 and +2 nucleosome occupancy levels, which lacked an NDR, also demonstrated the highest gene expression, suggesting that *S. purpuratus* may employ unique regulatory mechanisms that enable transcriptional access to promoters even in the absence of an NDR. While +1 nucleosomes are associated with poised gene expression and promoter proximal polymerase pausing in other organisms (Gilchrist et al., 2010; Jimeno-González et al., 2015), these nucleosomes are paired with a prominent NDR which is not observed in urchins. Our findings challenge the conventional idea that nucleosomes act as barriers to prevent transcriptional access to promoters (Abril-Garrido et al., 2023; Bai & Morozov, 2010; Han & Grunstein, 1988) and that nucleosome depleted regions are a defining feature of eukaryotic promoters (Jiang & Pugh, 2009; Mavrich et al., 2008b; Voong et al., 2016; Yuan et al., 2005). The observed sea urchin nucleosome distribution pattern necessitates a yet to be determined mechanism for engaging with, reading through, or rapidly remodeling the strongly positioned nucleosomes in the absence of an NDR. We further identified urchin promoters as being highly sensitive to MNase digestion. Within promoters, classes of genes defined by specific discrete sensitivity patterns are highly expressed. While there is some variation in sensitivity patterns across urchin sample replicates, the overall promoter sensitivity pattern was consistently dominated by highly positioned sensitive regions upstream of the TSS. Urchin sensitivity patterns are unique amongst eukaryotic organisms with discrete sensitive regions upstream of the TSS and few resistant regions. Notably, highly expressed urchin genes exhibit discrete MNase-sensitive regions just upstream or directly over the TSS, whereas highly expressed human genes are defined by highly positioned resistant regions. Our findings imply that urchin gene regulation mechanisms differ significantly from those of humans and other eukaryotes. Future studies focusing on chromatin structure and gene expression during urchin development or response to biotic or abiotic stressors are needed to fully understand the regulatory dynamics and mechanisms underpinning this relationship.

Collectively, our study elucidates the novel organization and regulatory characteristics of urchin chromatin architecture. Further research identifying the epigenetic patterns under an environmental stimulus (e.g. ocean acidification, marine heatwaves, starvation for marine organisms) can provide insight into ecological and climate change studies by providing methods to focus on the mechanistic explanation for differential gene expression. Chromatin architecture informs our understanding of epigenetic control of species under environmental stress and can delineate the molecular underpinnings of acclimation potential and plasticity. Although our work focuses on wild-caught *S. purpuratus*, the developed methodologies can be readily adapted to other organisms, tissue types, or conditions, offering a valuable framework for investigating gene regulatory mechanisms in echinoderms and other marine species.

## Methods

### Field Collections

Wild *Strongylocentrotus purpuratus* individuals were collected from Ucluelet, British Columbia, Canada (48.935716, -125.558527) in September 2021. Urchin gonads were dissected out of the animal using a sterile scalpel and crosslinked in 1% v/v formaldehyde solution in 34 parts per thousand salinity seawater for 10 minutes at room temperature. Gonad samples were flash frozen in liquid nitrogen (-200°C) and stored at -80°C. The animal used for MNase digestion was determined to be female based on SNP analysis of Nodal (LOC579010) and SOXE (LOC581729) promoters (as described by Pieplow et al., 2023).

### MNase cleavage and purification of nucleosomal DNA

Samples were thawed in 125mM Glycine to quench excess formaldehyde and the tissue was disrupted with a Dounce homogenizer. The homogenized tissue was divided into aliquots and pelleted by centrifugation for 10 minutes at 1000g at room temperature. The glycine supernatant was removed, and the sample was resuspended in nuclei isolation buffer (NIB: 10 mM HEPES at pH 7.8, 2 mM MgOAc2, 0.3 M sucrose, 1 mM CaCl2 and 1% Triton-X). The resuspended gonad nuclei were centrifuged for 10 minutes at 1000 x g at room temperature and pellets were rinsed twice in NIB. Rinsed and homogenized gonad samples diluted in NIB were pre-heated to 37°C prior to MNase digestion. A titration of predetermined MNase was added to each sample and pipetted gently to mix. MNase titrations were performed separately for each of the three replicates. Samples were incubated with 2 units of MNase at 37°C for 5-10 minutes for light digests or 15-20 minutes for heavy digests (Supp. Table 1). Reactions were halted with the addition of 0.5M EDTA and crosslinks were reversed by overnight incubation at 65°C in the presence of 0.05% sodium dodecyl sulfate (SDS) and 0.1mg Proteinase K. DNA from decrosslinked samples was isolated with silica columns and quantified using a nanodrop spectrophotometer. The size and quality of purified DNA was determined by running a 2% agarose gel in 1X TBE.

### Hamn THP-1 sample processing

THP-1 monocytes were obtained from the ATCC (TIB-202). Cells were cultured in RPMI-1640 media (Gibco) supplemented with 10% fetal bovine serum and 1% penicillin-streptomycin. Cells were maintained at 37°C with 5% CO2. Cells were crosslinked in 1% formaldehyde (Sigma-Aldrich) and incubated for 10 minutes at room temperature while rocking. Formaldehyde was quenched with 125mM glycine and cells were washed twice in PBS. Cells were resuspended in NIB and pelleted by centrifugation at 1000 x g for 10 minutes at 4°C. Nuclei were washed twice in NIB and nuclei purity was confirmed by DAPI staining. A concentration of 2.5×10^6^ nuclei were digested with 5U MNase for 10 minutes at 37°C alongside urchin samples as a control. Reactions were halted with 0.5M EDTA and nuclei were decrosslinked overnight at 65°C in the presence of 0.05% SDS and 0.1mg Proteinase K. DNA from decrosslinked samples was isolated with silica columns and quantified using a nanodrop spectrophotometer. Purified DNA from MNase digests was run on a 2% agarose gel in 1X TBE to evaluate fragment size and digest quality.

### DNA library preparation and sequencing

Mononucleosomal and subnucleosomal DNA fragments from triplicate urchin heavy and light digests were isolated using the Select-a-Size DNA Clean & Concentrator Kit (Zymo Research, cat D4080) by depletion of large fragments over 400bp. Between 200-300 ng of DNA was used to prepare libraries with the NEBNext Ultra II Library Prep Kit for Illumina with unique dual indices and 12 cycles of PCR amplification. Library size and quality was verified with a D1000 Tapestation. The molar concentration of each indexed library was determined by KAPA quantitative PCR and size corrected using sizing information from the Tapestation. Paired end 50 bp sequences were generated on the Florida State University College of Medicine’s NovaSeq 6000 Illumina sequencer.

### Raw file processing

The reads from heavy and light files were merged together to create total distribution files and processed in parallel with separate heavy and light digest files. All replicate files were processed in parallel. The quality of the raw fastq sequences was assessed with fastqc (v0.11.9) and trimmed with Trimmomatic v0.39 under moderate stringency conditions (Bolger et al., 2014). The post-trimming file quality was assessed with fastqc (v0.11.9) to ensure removal of poor-quality regions. The trimmed files were aligned with *bowtie2* (v2.3.5.1) to the Spur 5.0 genome assembly (Langmead & Salzberg, 2012). SAM files were converted to BAM format and low-quality reads were removed with *samtools* view 1.10 (Li et al., 2009). Sorted BAM files were indexed with *samtools* index and mitochondrial reads were removed with *samtools idxstats* and *samtools* view. Resulting BAM files were deduplicated with *picardtools markduplicates* (v2.27.3 Picard Toolkit, 2019). Samples were read depth normalized with *picardtools DownSampleSam* within each replicate for each heavy and light files and across replicates for total distribution files. The genome coverage was calculated with *Deeptools* (v3.5.5) *plotcoverage* and GC bias was assessed with *Deeptools computeGCBias* (Ramírez et al., 2014). Samples were compared with *Deeptools multibamsummary* and Pearson’s correlations were visualized with *plotCorrelation*.

### Nucleosome analysis

Nucleosome positioning was called with the *dpos* tool from *DANPOS3* (v2.2.2) for heavy, light, and merged samples with the -m (mate paired) option (Chen et al., 2013). Resulting wig files were converted to bigwig format with the *wigtoBigWig* tool from the UCSC genome browser (Kent et al., 2010). MNase sensitivity was determined with *Deeptools bigWigCompare* by calculating the log2ratio of light to heavy digests. Nucleosome distribution and sensitivity was plotted over genomic features with *Deeptools computeMatrix, plotProfile, and plotHeatmap*. Promoter sequences were isolated from the *S. purpuratus* ref seq gtf file from the UCSC genome browser (Clawson et al., 2023) with a custom python script. Genome wide and single loci views were generated with the UCSC genome browser (Raney et al., 2024).

### Public urchin data processing

Publicly available RNAseq data from adult *S. purpuratus* ovaries were downloaded from the NCBI SRA under accession SRR531958 (Tu et al., 2012, 2014). Files were trimmed with Trimmomatic (Bolger et al., 2014) and aligned to the Spur_5.0 genome assembly with HiSat2 (Kim et al., 2015). SAM files were converted to BAM format and low-quality reads were removed with *samtools* view 1.10 (Li et al., 2009). Stringtie (Pertea et al., 2015) was used to assemble and quantify transcripts. Expression data was intersected with gene names in the Deeptools kmeans clustering output in R (version 4.4.1) with custom R scripts to determine relationships between nucleosome distribution, sensitivity, and gene expression.

Publicly available PROseq data from developing *S. purpuratus* embryos were downloaded from the NCBI SRA under accessions GSE160460 and GSE234140 (Arenas-Mena et al., 2021; Arenas-Mena & Akin, 2023). Files were trimmed with Trimmomatic (Bolger et al., 2014) and aligned with Bowtie2 (Langmead & Salzberg, 2012) to the Spur_5.0 genome assembly. Deeptools (Ramírez et al., 2014) was used to call strand aware coverage profiles for each sample. Resulting coverage profiles were compared with promoter coordinates generated from the Spur_5.0 genome assembly to validate TSS coordinates.

Publicly available Pol II ChIPseq data from developing *S. purpuratus* embryos were downloaded from the NCBI SRA under accessions GSE160462 (Arenas-Mena et al., 2021). Files were trimmed with Trimmomatic (Bolger et al., 2014) and aligned with Bowtie2 (Langmead & Salzberg, 2012) to the Spur_5.0 genome assembly. SAM files were converted to BAM format and low-quality reads were removed with *samtools* view 1.10 (Li et al., 2009). Sorted BAM files were indexed with *samtools* index and mitochondrial reads were removed with *samtools idxstats* and *samtools* view. Resulting BAM files were deduplicated with *picardtools markduplicates* (v2.27.3 Picard Toolkit, 2019). Peaks were called with MACS3 v3.0.0b1 (Zhang et al., 2008) using the effective S.pur genome size. Deeptools (Ramírez et al., 2014) was used to generate heatmaps of called peaks for each sample.

### Public nucleosome data processing

MNase-seq data was obtained from the NCBI SRA for the following datasets: GSE254274, GSE26412, GSE78984, PRJNA297204, and GSE44269 (Mieczkowski et al., 2016; Parvathaneni et al., 2020; Xi et al., 2011; Zhang et al., 2014). Files were trimmed with Trimmomatic (Bolger et al., 2014) and aligned to respective reference genomes with Bowtie2: hs1/T2T, R64, MM10, BDGP6, B73, and GRCz11. SAM files were converted to BAM format and low-quality reads were removed with *samtools* view 1.10 (Li et al., 2009). Sorted BAM files were indexed with *samtools* index and mitochondrial reads were removed with *samtools idxstats* and *samtools* view. Resulting BAM files were deduplicated with *picardtools markduplicates* (v2.27.3 Picard Toolkit, 2019). Nucleosome positioning was called with the *dpos* tool from *DANPOS3* (v2.2.2) with the -m (mate paired) option for paired-end files and omitted for single end files (Chen et al., 2013). MNase sensitivity was determined with *Deeptools bigWigCompare* by calculating the log2ratio of light to heavy digests, when available. Nucleosome distribution and sensitivity was plotted over genomic features with *Deeptools computeMatrix, plotProfile, and plotHeatmap*. Genome wide and single loci views were generated with the UCSC genome browser (Raney et al., 2024). Promoter sequences were isolated from the appropriate ref seq gtf files on the UCSC genome browser (Clawson et al., 2023) with a custom python script.

## Data Access

All raw and processed urchin sequencing data generated in this study have been submitted to the NCBI Gene Expression Omnibus (GEO; https://www.ncbi.nlm.nih.gov/geo/) under accession number GSE283453. Source code used to process files is available as a supplementary file.

## Acknowledgements

Figure 1, the graphical abstract, and selected components of other figures were generated in Biorender.com. Silhouette images of each organism are by Chuanixn Yu (*Danio rerio*), Gemma Martínez-Redondo (*Sophophora pseudoobscura*), Harold N Eyster (*Strongylocentrotus purpuratus)*, Jiro Wada (*Mus musculus*), Katy Lawler (*Homo sapiens sapiens)*, Mason McNair (*Zea mays*), and Wayne Decatur (*Saccharomyces cerevisiae*). *© 2024. This work is openly licensed via CC BY 4.0 (*https://www.phylopic.org/permalinks/2a2ea455dea80ca4639ccfe9152cbd2bc84da69d0b6c45f29d0963ff4e23385e). Furthermore, we thank members of the Dennis and Okamoto labs for insightful conversations on results and data analysis.

## Author contributions

MJM and JMB contributed to conceptualization, methodology, formal analysis, data curation, and writing. JMB conducted figure visualization and software development. SEK contributed to formal analysis, data curation, and writing. LNA assisted with laboratory methodology and manuscript review. JHD and DKO contributed to funding acquisition, conceptualization, supervision, resources, and writing.

